# The *Achromobacter* Type 3 secretion system drives pyroptosis and immunopathology via independent activation of NLRC4 and NLRP3 inflammasomes

**DOI:** 10.1101/2023.04.06.535904

**Authors:** Keren Turton, Hannah Parks, Paulina Zarodkiewicz, Mohamad A. Hamad, Rachel Dwane, Georgiana Parau, Rebecca J. Ingram, Rebecca C. Coll, Clare E. Bryant, Miguel A. Valvano

## Abstract

*Achromobacter* species are newly recognized opportunistic, pro-inflammatory Gram-negative pathogens in immunocompromised individuals, but how they interact with the innate immune system to drive inflammation is poorly understood. We created *sctV* (Type 3 Secretion System baseplate) mutants in three *Achromobacter* clinical isolates from two species and showed that all three required the T3SS to induce cell death in human macrophages. Mutating other critical T3SS components also abolished cell death, which was restored by genetic complementation. Cell death of *Achromobacter*-infected macrophages was contact-dependent, enhanced by bacterial internalisation, and caused by inflammasome-dependent pyroptosis (typified by Gasdermin-D cleavage and IL-1β secretion). Macrophages deficient in the inflammasome sensors NLRC4 or NLRP3 underwent pyroptosis upon bacterial internalization but those deficient in both NLRC4 and NLRP3 did not, suggesting either sensor can mediate pyroptosis induction in a T3SS-dependent manner. Detailed analysis of the intracellular trafficking of one isolate indicated that the intracellular bacteria reside in an acidic LAMP-1/dextran-positive membrane compartment. Using an intranasal mouse infection model, we observed that *Achromobacter* damages lung structure and causes severe illness, contingent on a functional T3SS. Together, we demonstrate that *Achromobacter* species can survive phagocytosis by macrophages and promote macrophage cell death and inflammation by redundant mechanisms of pyroptosis induction.

## Introduction

Species of the genus *Achromobacter*, primarily found in soil, are emerging opportunistic Gram-negative pathogens in immunocompromised individuals^1^. In people with cystic fibrosis, the infection is recalcitrant, and re-colonisation even after lung transplant is common^2^. Disease severity and symptoms are comparable to those of *Pseudomonas aeruginosa*, a common cystic fibrosis pathogen^3^. Additionally, *Achromobacter* bacteraemia^4-6^ and infections in patients with hematologic and solid organ malignancies, renal failure, and various immunodeficiencies have been reported^1^. *Achromobacter* infections are very difficult to treat because multi-drug resistance is common, and in cases of chronic infections the bacteria frequently acquire further resistance over time^7,8^. This genus has received less attention than more ubiquitous lung pathogens, such as *Klebsiella* and *Pseudomonas*, but its increasing prevalence^9,10^ justifies further investigation. In addition to *A. xylosoxidans*, the primary focus in previous studies, several other *Achromobacter* species including *A. insuavis* and *A. ruhlandii* are clinically significant^11^.

*Achromobacter* species carry virulence and drug resistance genes that make them formidable during infection, particularly in immunocompromised individuals^11,12^. Previous studies have reported the biofilm potential of *A. xylosoxidans* and the presence of an uncharacterized heat-stable cytotoxin^9,12-15^. The importance of the *Achromobacter* Type 6 secretion system (T6SS) in interactions with co-infecting pathogens^16^ and the cytotoxicity of a particular Type 3 secretion system (T3SS) effector, AxoU, found in many strains of *A. xylosoxidans*^17^ have also been recently highlighted. Additionally, a repeats-in-toxin adhesion protein (RTX adhesin) contributes to the pathogenicity of *A. xylosoxidans* isolates^18^. The pro-inflammatory potential of *Achromobacter* infections in individuals with cystic fibrosis has been clinically established^19,20^, but how *Achromobacter* species interact with the innate immune system to elicit inflammation remains unknown. This work aims to unravel macrophage-*Achromobacter* interactions since macrophages are one of the front-line defenders of the immune system. We investigated macrophage cell death following *Achromobacter* infection using the *A. insuavis* isolate AC047 and two clinical isolates of *A. xylosoxidans*. AC047 is cytotoxic to human macrophages despite lacking the characterised virulence factors AxoU and RTX adhesin, suggesting that macrophage cell death mediated by *Achromobacter* species is multifactorial. In this study, we created a set of mutations in *A. insuavis* AC047 and two *A. xylosoxidans* isolates to delineate the role of T3SS components in infection and inflammation using human macrophage cell models and a mouse model of respiratory infection.

Pyroptosis is a form of pro-inflammatory programmed cell death that contributes to the recruitment of immune cells and can be induced by several Gram-negative pathogens (reviewed in ^21^). Accordingly, one can utilise attributes that *Achromobacter* spp. have in common with other Gram-negative pathogens to predict triggers of pyroptosis induction. Several genera, like *Burkholderia*, induce pyroptosis via several different inflammasome sensor pathways (reviewed in ^22^). In this work, we show that human macrophages undergo pyroptosis when infected by *Achromobacter* spp. and that this process is T3SS-dependent in multiple clinical strains. We used a panel of human macrophage cell lines deficient in specific inflammasome components to assess the contributions of Nod-like receptor (NLR) family CARD domain-containing protein 4 (NLRC4) and NLR family pyrin domain containing 3 (NLRP3), which are sensors of pathogen-associated molecular patterns^21^. We demonstrate that while the NLRC4 and NLRP3 inflammasome sensors are each sufficient for pyroptosis induction, neither is absolutely required. Additionally, we utilised a less cytotoxic *A. xylosoxidans* strain to study the intracellular trafficking of the bacterium in macrophages, revealing that intracellular survival of *Achromobacter* occurs in a late phagosome and is T3SS-independent. The T3SS was also required to elicit lung inflammation and tissue damage in a mouse infection model, indicating that this system plays a significant role in the pathogenicity of *Achromobacter in vivo*.

## Results

### The T3SS of *Achromobacter* species is necessary and sufficient to induce macrophage cell death

The ability of the *Achromobacter* clinical isolates *A. insuavis* AC047 and *A. xylosoxidans* QV306 to induce cell death in macrophages was assessed by monitoring lactate dehydrogenase (LDH) release. Both strains induced cell death in THP-1 macrophage-like cells, albeit on different timescales **(Figure 1A)**, suggesting the extent of cytotoxicity is strain-dependent. Live but not heat-killed bacteria induced cytotoxicity. To investigate whether cytotoxicity by *Achromobacter* species is associated with the T3SS, we constructed mutants in the *sctV* gene encoding the baseplate protein of the T3SS in AC047, QV306 and in another *A. xylosoxidans* isolate, AC055. This gene was selected for our initial mutagenesis experiments since previous work in *Salmonella*^23^ and *Yersinia*^24^ showed that SctV functions in T3SS secretion and T3SS needle assembly. We generated an unmarked *sctV* deletion in AC047 and insertional *sctV* mutants in *A. xylosoxidans* QV306 and AC055. All three mutants lost the ability to cause macrophage cell death **(Figure 1B)**, indicating that the *Achromobacter* T3SS is required for cell death induction in infected THP-1 macrophage-like cells.

**Figure 1.**
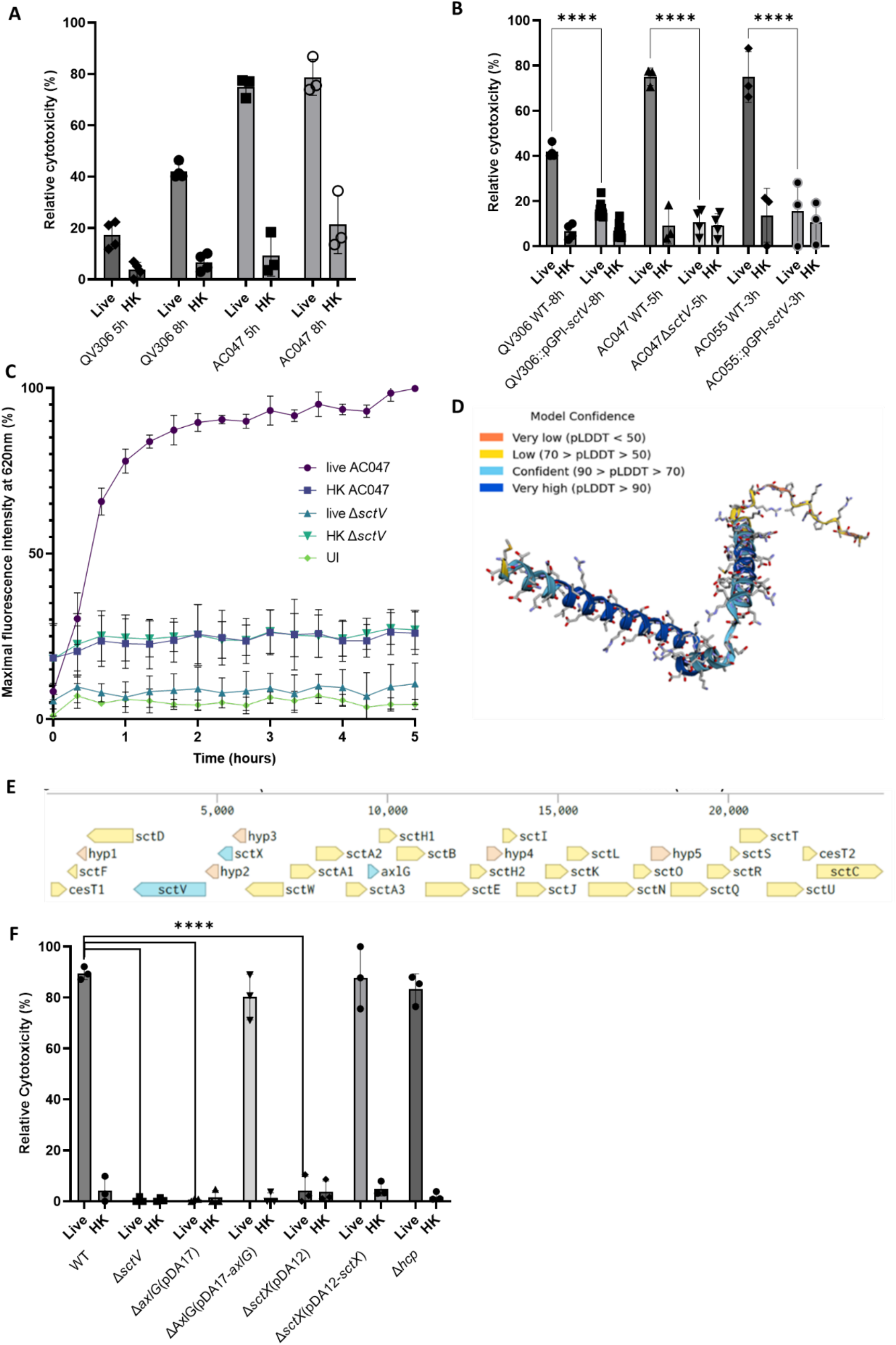
The T3SS is required for cytotoxicity induction in macrophages. **A**. LDH release of THP-1 cells infected with live and heat-killed (HK) bacteria **B.** LDH release of THP-1 cells infected with live and *sctV* mutants **C.** Propidium iodide uptake time-course assay of HMDM infection with strain AC047 and Δ*sctV* mutant. **D.** Alphafold prediction of AC047’s AxlG structure. **E.** Schematic of AC047 T3SS operon generated using Benchling. **F.** LDH release of HMDM infected with live and heat-killed (HK) AC047 and its Δ*sctX* and Δ*axlG* mutants at 5 hours p.i. Data represent at least three biological replicates (four technical each), UI = uninfected, MOI 20. **** signifies p<0.0001 by t-test, error bars SD.

We also examined primary human monocyte-derived macrophages (HMDMs) infected with AC047 using a propidium iodide (PI) uptake assay to follow the time-course of cytotoxicity. As observed for LDH release in THP-1 macrophages, wildtype but not AC047Δ*sctV*, induced rapid cytotoxicity in HMDMs (**Figure 1C**), reaching nearly 80% of its peak within 1 hour. In contrast, heat-killed or *sctV*-deficient bacteria did not induce significant PI uptake. Macrophages exposed to heat-killed bacteria showed basal propidium iodide readings around 10% higher than uninfected macrophages, and macrophages infected with live Δ*sctV* (**Figure 1C**). We attributed these slightly higher values to PI fluorescence arising from the bacterial DNA in the heat-killed bacterial controls, since PI cannot access live bacterial cells.

To verify that an intact T3SS was necessary for cytotoxicity in HMDMs, we attempted to complement the *sctV* gene deletion in AC047, but despite several trials, complementation was unsuccessful possibly due to either polar effects of the mutations or instability of the expressed SctV protein. Therefore, we created unmarked deletions of two other genes, *sctX* (a homologue of *yscX*) and *axlG*. In *Yersinia*, YscX is essential in early secretion from the T3SS^25,26^. The AxlG protein, in contrast, consists of fewer than 120 amino acids with a predicted pI of 4.81 and several α-helices predicted by AlphaFold^27^, which form a V-shape fold (**Figure 1D**), all attributes of a T3SS chaperone^28^. Analysis of the T3SS cluster by cblaster in 208 *Achromobacter* genomes revealed that *axlG* was immediately adjacent to *sctA* in 98% of them^29^. The standard nomenclature was used to refer to the T3SS genes whenever possible^30^, but *axlG* is not a core component of the T3SS in other bacterial species. We therefore named this gene according to the label system created by Pickrum *et al*. for the equivalent gene in their *Achromobacter* strain^17^ (schematic of AC047 operon in **Figure 1E**). Knockout of *sctX* and *axlG* genes abrogated cytotoxicity, but complementation with the respective parental genes restored it (**Figure 1F**). Many *Achromobacter* species have a Type VI secretion system (T6SS), which can play a role in the infection of respiratory epithelial cells^16^. To rule out the T6SS as a contributor to macrophage cell death, we generated a mutant lacking Hcp, an essential protein component of the T6SS apparatus, with only one copy of the gene in AC047. The disruption of *hcp* does not affect LDH release (**Figure 1F**); showing that infection-induced cell death in macrophages is T6SS-independent. Moreover, there was no evidence of residual cytotoxicity found in macrophages infected with the T3SS-defective mutants where the T6SS remained intact.

### Cell death is contact-dependent, and enhanced by bacterial internalisation

Mantovani *et al*.^15^ suggested that some *A. xylosoxidans* strains could secrete an uncharacterized heat-stable cytotoxic factor. Therefore, we investigated whether T3SS effectors secreted into the media could cause cytotoxicity in macrophages. Because *Achromobacter* species are multidrug-resistant, we could not perform a classical gentamicin assay to kill extracellular bacteria during infection. Various other combinations of antibiotics were trialled but ultimately deemed unsatisfactory since the high antibiotic thresholds required led to macrophage cytotoxicity, or to killing of intracellular bacteria as infection-induced cytotoxicity culminated in membrane disruption of host macrophages. Although we washed bacteria repeatedly before suspension in RPMI to remove effectors in the culture media, it was possible that bacteria could secrete more effectors during infection. To circumnavigate these issues, we used transwells^31^ to demonstrate whether bacteria-cell contact was required to induce cytotoxicity. As shown in **Figure S1A**, *Achromobacter* must be in contact with macrophages to cause cytotoxicity as LDH release is abrogated in the transwells.

We next investigated whether internalisation was a pre-requisite for cytotoxicity by pre-treating macrophages with 5 μg/ml cytochalasin D (CD), which inhibits phagocytosis^32^. In HMDMs CD delayed the onset of cytotoxicity as measured by the PI permeability assay (**Figure S1B**). Fluorescence did not increase in the uninfected cytochalasin D-treated control or in macrophages exposed to the DMSO vehicle. We also examined the effect of CD in THP-1 macrophages infected with QV306, AC047 and AC055 using confocal microscopy by following bacteria expressing the mCherry fluorescent protein. In **Figure S1C** at 5 hours post infection (p.i.), we observed no internalised *Achromobacter* species inside CD-treated cells. However, the *A. xylosoxidans* QV306 and AC055 bacterial cells adhered to the outside of CD-treated cells, while the *A. insuavis* AC047 bacteria did not. We speculate these differences reflect the presence of the RTX adhesin^18^, which is encoded in *A. xylosoxidans* QV306 and AC055, but absent in the *A. insuavis* AC047 genome, and can promote cytotoxicity upon bacterial contact with the surface of target cells. Together, these experiments demonstrate that macrophage cell death requires both bacteria-cell contact and internalisation.

### *Achromobacter* species reside in a late phagosome compartment in a T3SS-independent manner

*A. xylosoxidans* has been reported to survive intracellularly without significant replication, exiting infected cells upon cytotoxic membrane rupture^18^. To understand the intracellular trafficking and survival strategy of *Achromobacter* species after phagocytosis, we investigated the maturation of the *Achromobacter* containing vacuole (AcV) in THP-1 macrophages infected with the less acutely cytotoxic *A. xylosoxidans* strain QV306 expressing mCherry. After phagocytosis, the AcV transiently acquired the early endosomal antigen 1 (EEA1) marker, but its association with this marker declined over 1-hour p.i. (**Figure 2A**). In contrast, over time the AcV increasingly associated with the late endosome/phagolysosome marker LAMP-1 **(Figure 2B)**. Image quantifications **(Figure 2C and D)** demonstrate that while the association of the AcVs with EEA1 decreases over time, the association with LAMP-1 increases and virtually all AcVs are LAMP-1-positive after 3 hours p.i. Similar kinetics of the AcV localising with LAMP-1 are observed in HMDMs **(Figure S2).**

**Figure 2.**
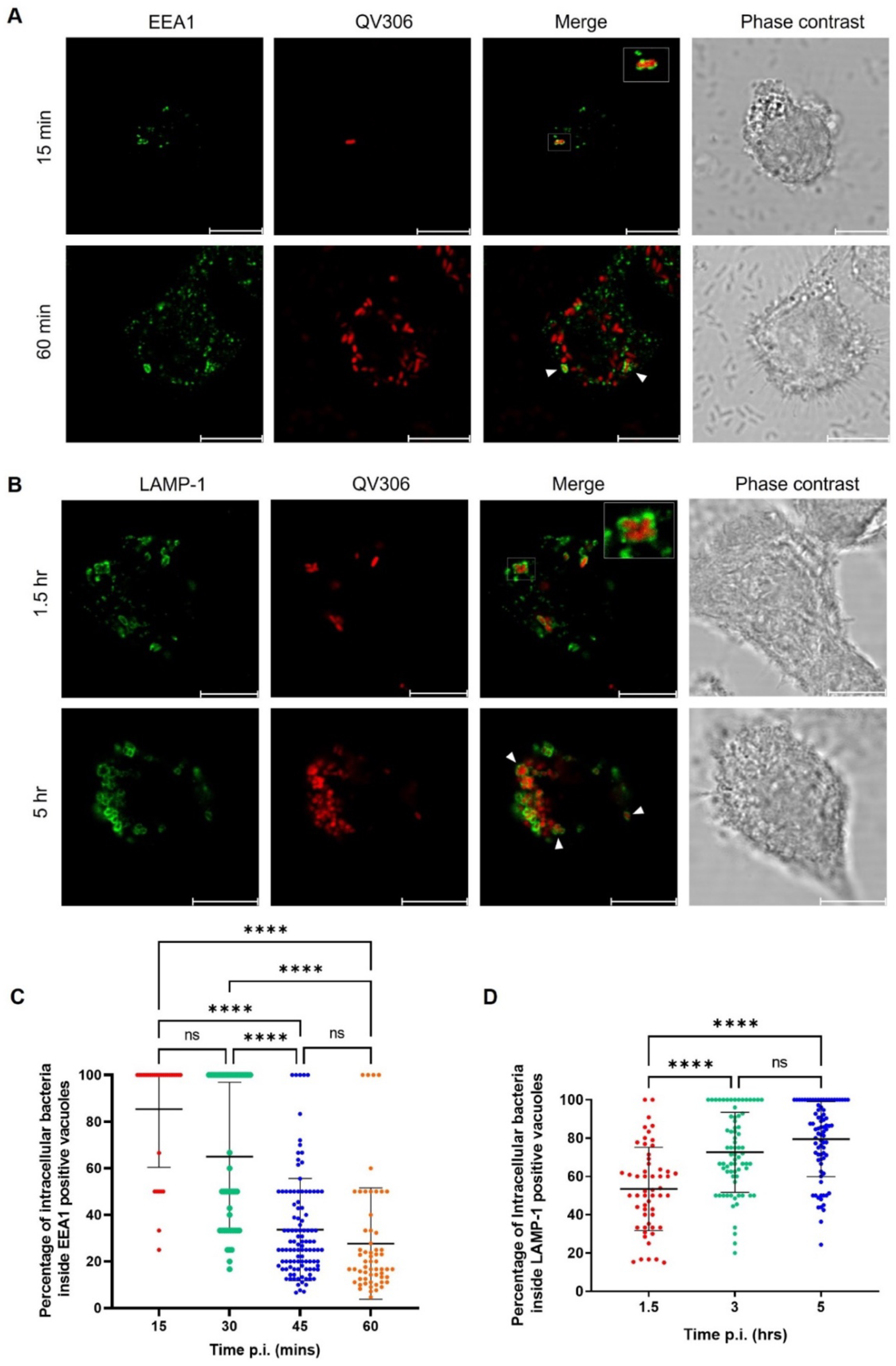
*A. xylosoxidans* QV306 infecting THP-1 macrophages resides in a vacuole that resembles a late endosomal compartment. **A.** macrophages at 30- and 60-minutes p.i. with live QV306 bacteria colocalising with EEA1. **B.** macrophages 1.5 and 5 hrs p.i. with live *A. xylosoxidans* colocalising with LAMP-1. Images taken with × 63 magnification on Leica SP8 confocal microscope. MOI = 80. Scale bar = 10 µm. **C.** The percentage of intracellular bacteria present in EEA1 positive vacuoles was assessed in 40-100 macrophages. **D.** The percentage of intracellular bacteria present in LAMP-1-positive vacuoles was assessed in 80 macrophages. **** p < 0.0001 by Kruskal-Wallis test. Error bars represent standard deviation from the mean; datasets from three biological replicates.

Other pathogens associated with cystic fibrosis, such as *Burkholderia cenocepacia,* accumulate the autophagy marker LC3b early after phagocytosis^33^. In contrast to THP-1 macrophages infected with *B. cenocepacia*, which showed the presence of LC3b associated with the bacteria-containing vacuole at 3 hours p.i., the AcV was not associated with LC3b (**Figure S3**), indicating that the AcV is not an autophagic vacuole. Dextran fluorescein is a fluid phase marker that demonstrates co-localisation in the phagolysosome, where ingested bacteria are degraded. At 3- and 5-hours p.i., the AcV co-localises with dextran fluorescein **(Figure S4)**. This supports our LAMP-1 co-localisation data **(Figure 2B)**, corroborating that the bacteria are in a late phagosome compartment. We also investigated whether the AcV matures to become acidic by evaluating the accumulation of Lysotracker green DND. The results show that some but not all AcVs appear to be acidic compartments **(Figure S5)**. This heterogeneity could be due to the presence of extracellular bacteria that are differentially internalised over time during the experiment since we cannot effectively kill either the residual extracellular bacteria at the onset of the experiment or intracellular bacteria released after macrophage cell lysis. We also infected HMDMs with mCherry-labelled AC047Δ*sctV* to investigate whether the T3SS is required for intracellular survival. The results show that the mutant can be internalised in the absence of a T3SS and accumulates in a LAMP-1-positive vacuole containing intact bacterial cells for at least 24 hours p.i. (**Figure S6**). Together, our data indicate that *Achromobacter* strains can reside in a late phagosome compartment and this process is T3SS independent.

### *Achromobacter*-induced macrophage cell death occurs by T3SS-dependent pyroptosis and requires NLRP3 and NLRC4 inflammasome sensors

The innate immune system can recognise common PAMPs of bacterial pathogens, like lipopolysaccharide (LPS) and T3SS components, triggering pyroptosis^21^. To establish whether macrophage cell death is due to pyroptosis, we examined cell lysates from THP-1 macrophages infected with several clinical isolates of *Achromobacter* for Gasdermin-D (GSDMD) cleavage, a proxy for a pro-inflammatory response leading to the formation of membrane pores and release of mature interleukin(IL)-1β/IL-18 into the supernatant^34^. GSDMD cleavage (by N-terminus blotting) was observed in all live but not heat killed strains containing T3SS (AC035, AC055, AC088) to a level comparable with GSDMD cleavage triggered by NLRP3 activation (LPS and nigericin) (**Figure 3A**). The *A. xylosoxidans* isolates AC055 and AC088, and the *Achromobacter* species cluster-II isolate AC035, but not the *A. xylosoxidans* AC011 and ATCC27061 (type strain) isolates contained T3SS systems, based on DNA sequencing and PCR results **(Figure S7)**. An intact T3SS is required for pyroptosis induction by AC047 and AC055 as GSDMD cleavage was absent in HMDM infected with *sctV* mutants (**Figure 3B**), but unaffected in the T6SS (*hcp*) mutant. As expected, complementing the deleted *axlG* gene in the AC047Δ*axlG* mutant restored GSDMD cleavage (**Figure 3C**). Live AC047 infection resulted in IL-1β release, an effect not seen in the T3SS-deficient mutant (**Figure 3D**). When complemented, AC047Δ*axlG* released IL-1β levels comparable to wildtype mirroring the restoration of GSDMD cleavage. Together, these data indicate that a functional T3SS is required to induce pyroptosis in THP-1 macrophages and HMDMs.

**Figure 3.**
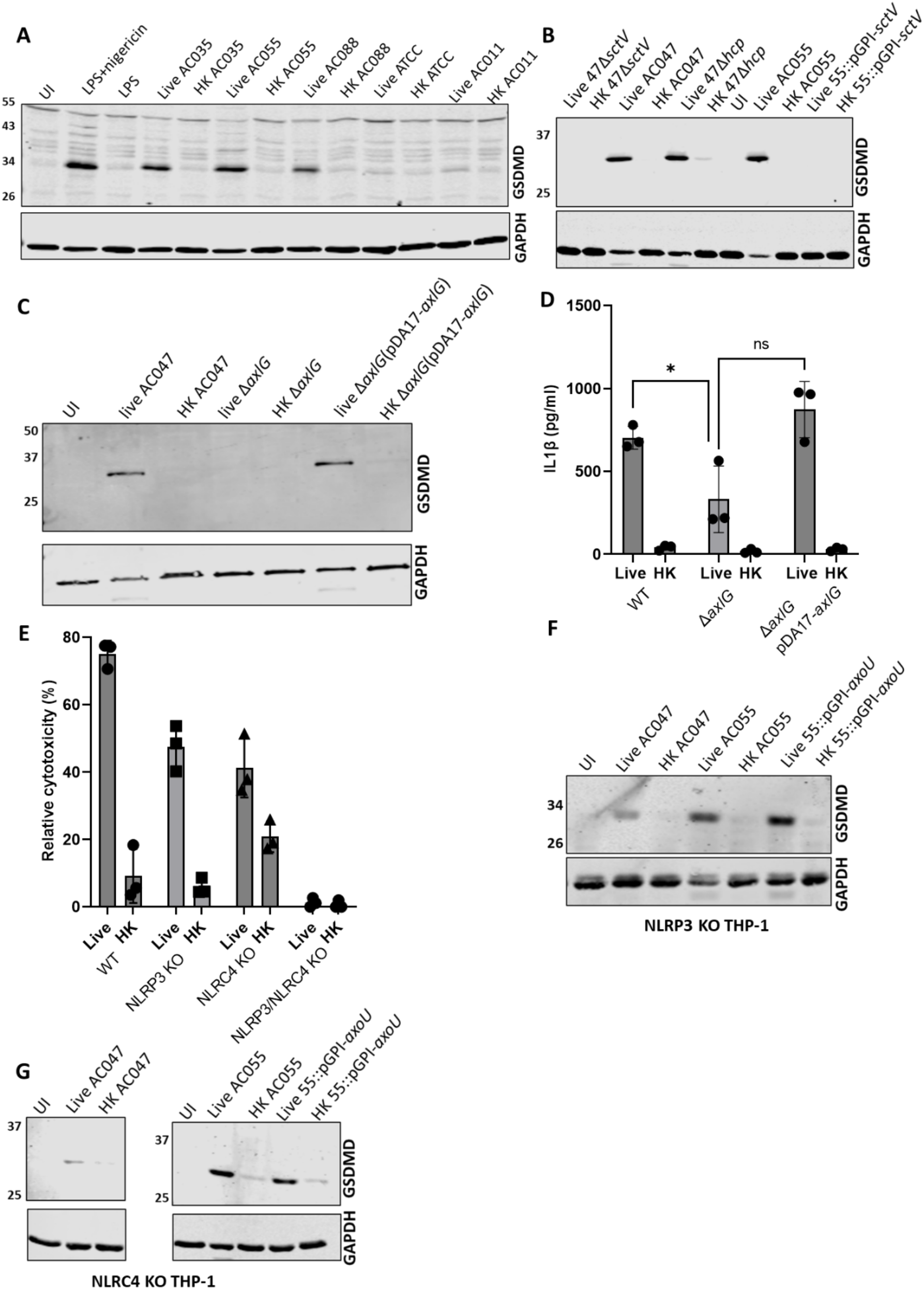
The Type Three Secretion System is required for NLRC4- and NLRP3-dependent pyroptosis in macrophages. **A.** Gasdermin-D cleavage in THP-1 infected with live and heat-killed (HK) *Achromobacter* spp, antibody CST #93709. **B-C.** Gasdermin-D cleavage in HMDM infected with live and HK AC047, AC055 and mutants. Antibody (CST #36425) detects N-terminus of cleaved GSDMD. **D.** ELISA of IL-1β in supernatants of infected HMDM (3h p.i.). Data represents three biological replicates, * signifies p<0.05 by t-test, error bars SD MOI 20. **E.** LDH release of infected KO THP-1 lines. Gasdermin-D and GAPDH immunoblots of NLRP3 KO **(F)** and NLRC4 KO **(G)** THP-1 infected with live and HK bacteria. MOI 20, infection 5 hours, error bars SD. Data represents three biological replicates. Antibody (CST #36425) detects N-terminus of cleaved GSDMD.

Two inflammasome sensors, NLRC4 and NLRP3, are known to detect intracellular Gram-negative pathogens^21^. Therefore, our next set of experiments focused on establishing whether they play a role in detecting *Achromobacter* species. In particular, NLR family of apoptosis inhibitory proteins (NAIPs) are innate immune sensors^35^ that can co-assemble with NLRC4 upon detection of T3SS components^36^ and in some instances, depending on the macrophage model, NLRP3 can also be involved^37^. We investigated whether NLRC4 or NLRP3 participate in detecting *Achromobacter* infection using a set of NLRP3, NLRC4, and NLRP3/NLRC4 knockout THP-1 cells. The results show that NLRC4 and NLRP3 sensors were each sufficient, but neither was solely necessary for cytotoxicity (**Figure 3E**), which was corroborated by immunoblotting data indicating that pyroptosis still occurred in each of the single KO cell lines (**Figure 3F-G**). Flagellin is involved in NLRP3-dependent pyroptosis induction in *Salmonella* infection^37^. We constructed a *fliC* mutant in AC047 to determine if the absence of *fliC* affected cytotoxicity in AC047 infection of NLRC4 KO THP-1. There was no significant difference between WT and Δ*fliC* (**Figure S8**), suggesting an alternate mechanism.

### Effector AxoU is cytotoxic independently of NLRC4 and NLRP3, and is not required for pyroptosis

Previous work in *A. xylosoxidans* identified AxoU, a homolog of the *Pseudomonas aeruginosa* T3SS-secreted ExoU phospholipase A_2_, which induced cell death in macrophages when expressed in *P. aeruginosa* and delivered by the *P. aeruginosa*’s T3SS^17^. To investigate the role of AxoU in cell death, we generated mutants of AC055 with disrupted *axoU* and *sctV* genes. As shown in **Figure S9A**, AC055 is still cytotoxic for NLRC4/NLRP3 KO THP-1 while disrupting *axoU* abrogates cell death. Disrupting *sctV* prevented cell death in HMDM and THP-1 (**Figure 1B**) but disrupting *axoU* alone did not, demonstrating that AxoU is not the only cytotoxic T3SS effector (**Figure S9B**). Wildtype AC055, unlike the AC055 *axoU* mutant, was cytotoxic to NLRC4/NLRP3 KO THP-1 (**Figure S9A**), showing that *axoU*-mediated cytotoxicity is independent of these sensors. Similarly, treatment with pan-caspase inhibitor Z-Val-Ala-Asp(OMe)-fluoromethylketone (Z-VAD-FMK) abrogated cell death induction by the AC055 mutant lacking AxoU (**Figure S9C**), but not by wildtype AC055 with a functional AxoU. Z-VAD-FMK inhibits both pyroptosis and apoptosis, suggesting AxoU induces cytotoxicity independently of both these pathways. Because AxoU could have redundant means of inducing toxicity, we assessed whether AC055::pGPI-AxoU could still elicit GSDMD cleavage. In NLRC4 KO, NLRP3 KO (**Figure 3F-G**) and WT THP-1 cells (**Figure S9D**), GSDMD cleavage occurred even when the *axoU* gene was disrupted, while in the NLRC4/NLRP3 KO THP-1 no pyroptosis occurred upon infection with wildtype or AC055::pGPI-*axoU* (**Figure S9D)**. The *axoU* gene is present in many *A. xylosoxidans* strains but absent in the *A. insuavis* AC047, which can induce pyroptosis. AC055::pGPI-AxoU mirrors the phenotypes observed for AC047 (**Figures 3E, Figure S9B-C**). Together, these data indicate AxoU is not necessary for pyroptosis induction.

### *Achromobacter* T3SS is required for virulence and inflammation *in vivo*

We also investigated the pathogenic role of the T3SS in two *in vivo* models. We first compared AC047 wildtype and AC047Δ*sctV* in the *Galleria mellonella* wax moth model^38^. Larvae injected with wildtype AC047 died within 5 days p.i., whereas survival of larvae injected with AC047Δ*sctV* was comparable to the PBS-injected control (**Figure S10**), demonstrating that that a functional T3SS is required for *Achromobacter* virulence *in vivo*. To assess whether the T3SS is required for *Achromobacter* to establish a respiratory infection, we inoculated C57BL/6 mice intranasally. Mice administered 10^9^ and 10^8^ CFU wildtype AC047 died within 6 hours post-inoculation, whereas mice treated with the same dose of AC047Δ*sctV* showed no symptoms (**Figure 4A**). Inoculation with 10^7^ CFU of wild-type AC047 resulted in the recovery of bacteria from the lung but there was no development of symptoms of illness, as seen with higher doses (**Figure 4B**). The recovered bacteria showed a reduction in CFU/lung with respect to the initial inoculum, suggesting they can be cleared over time. However, an induction of inflammatory response in the lung was observed based on increased IL-6 production, which plays a role in the acute phase response (**Figure 4C)**. Other cell types rely on IL-1β-dependent IL-6 synthesis/release^39,40^, so we speculate that inflammasome activation influences IL-6 production in our model. Infection with 10^7^ CFU wild-type and AC047Δ*sctV* shows that a lack of T3SS results in slower bacterial clearance from the lungs and significantly lower inflammatory responses in comparison to wildtype, as seen by reduced IL-6 and keratinocyte chemoattractant (KC) levels (**Figure 4D-F**). These data suggest that lack of T3SS may be responsible for a reduced inflammatory response *in vivo*, possibly due to inability of AC047Δ*sctV* to activate NAIP/NLRC4 and NLRP3 innate immune responses, which also leads to decreased efficiency of pathogen clearance. Supporting this idea, lung histology demonstrated that the AC047Δ*sctV*-infected mice had no obvious signs of inflammation or lung damage (**Figure 4G**). However, wildtype-infected mice showed >4-fold increase in alveolar septa thickness and neutrophils in the interstitial space, which are parameters for assessment of acute lung injury^39^. We observed more alveolar wall damage (mean linear intercept) in the AC047-infected mice in comparison to the AC047Δ*sctV*-infected mice. Together, our data suggest that the T3SS contributes to eliciting an inflammatory response and reduction of bacterial load in the lung, but which causes significant tissue damage, while the absence of T3SS-mediated tissue inflammation leads to reduced clearance from the lung.

**Figure 4.**
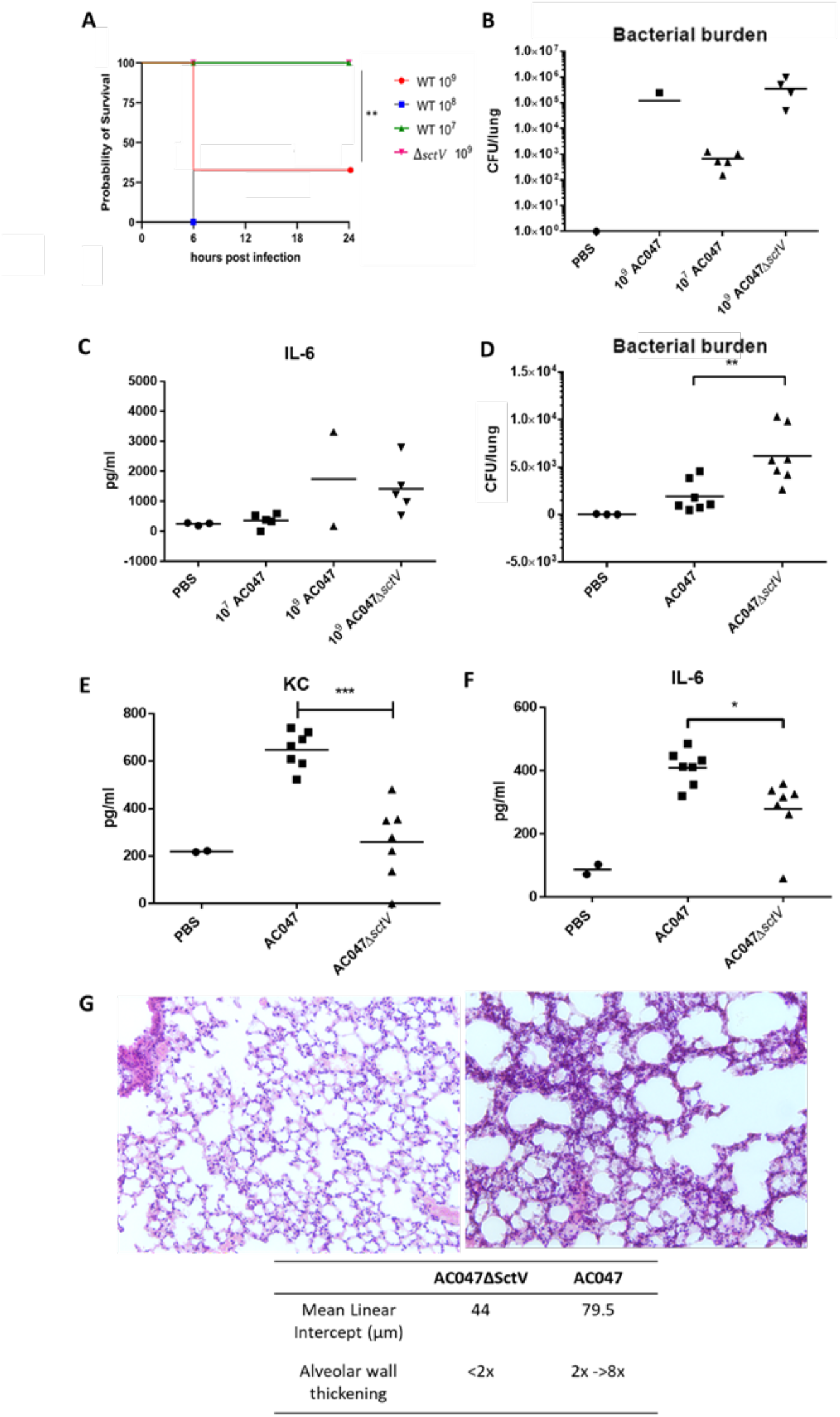
AC047Δ*sctV* is less virulent, causes lower inflammatory response and does not cause acute lung damage in comparison to AC047 in mice. **A,B,C.** Mice infected intranasally with 10^9^ and 10^8^ of AC047 succumb within 6 hours post infection. In comparison mice infected with 10^9^ of AC047Δ*sctV* survive 24 hours without developing any symptoms but produce an inflammatory response shown by increased IL-6. **D,E,F.** Mice infected intranasally with 10^7^ of AC047Δ*sctV* have lower recoverable bacterial load from lung homogenates, and show a lower inflammatory response shown by IL-6 and KC levels. **G.** AC047Δ*sctV* infected mice (left) show smaller alveolar destruction and no wall thickening, in comparison to AC047 infected mice (right). Scale bar = 50μm. * signifies p<0.05, ** p<0.005 and *** p<0.001 by t-test (ELISAs) or Mantel-Cox test (survival).

## Discussion

This study reveals how clinical isolates of *A. insuavis* and *A. xylosoxidans* exploit the T3SS to interact with human macrophages in a proinflammatory fashion resulting in pyroptosis. The T3SS, a bacterial nanomachine that delivers protein effectors into eukaryotic cells, is a crucial mediator of host-microbe interactions for many Gram-negative pathogens^41^. We established that the T3SS-defective *Achromobacter* mutants lost the ability to induce pyroptosis in macrophages, which required bacteria-macrophage contact and was enhanced by internalisation. When evaluating other clinical isolates in our collection, including the type-strain for the genus *Achromobacter* (ATCC27061), it became clear that those capable of killing macrophages also had T3SS operons, suggesting T3SS-mediated pyroptosis could be of broad clinical relevance.

The high-level multidrug antibiotic resistance of *Achromobacter* species, particularly of *A. xylosoxidans*, hampers the genetic manipulation of this genus due to the lack of appropriate drug selection markers. We discovered that the *A. insuavis* clinical strain AC047 was genetically tractable because it lacked the tetracycline resistance commonly found in *A. xylosoxidans*^11^, which enabled us to use a two-step recombinational approach leading to the generation of markerless deletion mutants by adapting a mutagenesis protocol previously developed for *Burkholderia*^42^. This approach allowed us to create a set of deletion mutants that helped establish unequivocally the T3SS’s role in *Achromobacter* interactions with human macrophages and in infections in animal models.

The clinical isolates of *A. insuavis* and *A. xylosoxidans* investigated here induced macrophage cell death, but the extent of cytotoxicity was strain dependent. Of the three strains examined, cytotoxicity was rapid and extensive in *A. xylosoxidans* AC055, while *A. xylosoxidans* QV306 was the least toxic and *A. insuavis* AC047 displayed intermediate toxicity. These differences could merit further study, given that both *A. xylosoxidans* strains carry intact *sacUaxoU* operons encoding the AxoU phospholipase and its cognate chaperone, and the T3SS gene clusters are highly conserved in all three strains. The mechanism of cell death was attributed to pyroptosis, based on the hallmark cleavage of GSDMD and the release of IL-1β. T3SS effectors could contribute to modulation or evasion of pyroptosis, as this is a known phenomenon in other Gram-negative bacteria, like YopK in *Yersinia*^43^. Bacteria can employ other mechanisms for immune evasion^44,45^, and it is possible that rather than AC055 having a particularly unique virulence effector, QV306 may have a T3SS effector that enables delayed pyroptosis. Alternatively, the T3SS regulation may differ among the three strains, which could account for inter-strain variation in the degree of pyroptotic cell death.

The relatively low cytotoxicity of *A. xylosoxidans* QV306 made this strain useful to study the intracellular lifestyle of *Achromobacter*. After establishing that T3SS deletion did not affect internalisation, we visualised internalised bacteria by immunostaining, demonstrating that they can survive for extended periods (at least up to 24 hours) in a membrane-bound *Achromobacter-* containing vacuole (AcV). Experiments with specific membrane and fluid-phase fluorescent markers revealed that the AcV matures from an early phagosome into a late phagolysosome that is partially acidified, suggesting intracellular *Achromobacter* may interfere with the normal acidification process associated to the phagolysosome maturation pathway. Recapitulation of these experiments with the T3SS-defective *A. insuavis* AC047Δ*sctV* demonstrated this secretion system is not required for intracellular survival, indicating that other bacterial components are involved in the intracellular lifestyle of *Achromobacter* species.

The *Achromobacter* strains induced pyroptosis by engaging either the NLRC4 or NLRP3 sensors, as demonstrated by infections in THP-1 cells with single and double knockouts in these sensors. However, our experiments could not distinguish whether effectors or structural T3SS components were detected by the inflammasome sensors. Detection of structural proteins of the assembled T3SS needle by NAIP/NLRC4-mediated sensing is a known phenomenon in *Salmonella, Shigella, Burkholderia* and other bacteria^36,46,47^. Because the *Achromobacter* isolates induced pyroptosis in NLRP3 KO THP-1 cells but not in the NLRC4/NLRP3 double KO cells, we conclude that the NLRC4 pathway is sufficient for pyroptosis induction, possibly by recognizing structural components of the T3SS that remain to be elucidated. However, the observation that NLRC4 was dispensable for T3SS-mediated pyroptosis in the presence of a functional NLRP3 was intriguing. It was recently demonstrated that in *Salmonella-*infected THP-1 cells, the bacterial flagellin activates NLRP3 while the PrgI(SctF) T3SS structural component activates NLRC4^37^. The absence of flagellin did not affect AC047’s ability to induce cytotoxicity in NLRC4 KO THP-1 (**Figure S7**), ruling out this potential mechanism in our model.

This study also demonstrated that AxoU can induce cell death independently of NLRC4 and NLRP3, and is not necessary for pyroptosis, since in its absence pyroptosis can still occur via either sensor. AxoU-induced cytotoxicity occurs even in the presence of a pan-caspase inhibitor, which abrogates cell death of macrophages infected with strains lacking AxoU. Moreover, *Bordetella bronchiseptica* (an evolutionary close relative of *Achromobacter*) can induce T3SS-dependent but caspase-1-independent necrosis^48^. Induction of pyroptosis by AxoU via a pathway independent of NLRP3 or NLRC4 is unlikely since *A. xylosoxidans* AC055, which carries an intact *axoU* gene, failed to induce pyroptosis in the NLRC4/NLRP3 KO THP-1, suggesting that AxoU is probably causing necrosis. This conclusion is also consistent with the function of the ExoU, the AxoU homologue in *P. aeruginosa*, which is cytotoxic to macrophages but inhibitory of caspase-1 activity^49^.

Our *in vivo* infection models also showed that T3SS-deficient *A. insuavis* was much less virulent than wildtype in systemic infection of *G. mellonella* larvae and in murine lung infections. Moreover, proinflammatory lung tissue damage occurred in mice infected with wildtype but not in those infected with the T3SS-deficient mutant. These findings agree with a recent study showing that *A. xylosoxidans* induced acute lung inflammation in mice^50^, which is consistent with the idea that macrophages undergo pyroptosis, a highly proinflammatory event. T3SS-defective bacteria were less efficiently cleared than wildtype, suggesting the T3SS may have a role in alerting the immune system, ultimately leading to bacterial clearance. A similar role was previously established for the T6SS secreted effector TecA, a bacterial deamidase that activates the pyrin inflammasome^51^.

In summary, this study shows that *Achromobacter* species can interact with human macrophages to induce pyroptosis by engaging NLRC4 or NLRP3 sensors in a process that is T3SS-dependent. We also show that pyroptosis is AxoU-independent and does not require the RTX adhesin, both of which are only present in *A. xylosoxidans* isolates^17,18^, demonstrating that the T3SS is a common denominator of pathogenicity in *Achromobacter* species. We propose that *Achromobacter* interactions with innate immune cells are particularly critical to determine infection and inflammation under conditions where the host is partially immunosuppressed, such as in individual with cystic fibrosis or other conditions, where *Achromobacter* species find a favourable niche to colonize and infect.

## Materials and Methods

### Cell culture

Wildtype THP-1 cells, and THP-1 with CRISPR-based knockouts of NLRP3^52^, NLRC4, and NLRC4/NLRP3^37^ were each cultured in RPMI 1640 supplemented with 10% heat-inactivated FBS and 1% penicillin-streptomycin. Cells were seeded at a density of 2 x 10^5^ cells/ml and differentiated in 80 ng/ml PMA for 3 days, then rested overnight without PMA before infection. On the day of infection, cells were refreshed with antibiotic-free media at least an hour beforehand.

### HMDM preparation

Buffy coats were prepared by Ficoll-Paque gradient density fractionation, and positively selected by incubation with CD14+ beads. Isolated cells were cultured in IMDM 10% FBS and 1% penicillin-streptomycin 50 ng/ml GM-CSF for seven days. Cells were plated at a density of 5 x 10^5^ cells/ml with GM-CSF and infected the next day. On the day of infection, cells were refreshed with antibiotic-free media an hour before infection.

### Strains

Clinical strains used in this work were part of a collection of *Achromobacter* species sputum isolates originated from individuals with cystic fibrosis, which were obtained from the Antimicrobial Resistance and Healthcare Associated Infections Reference Unit of the UK Health Security Agency. Strains were grown in a semi-defined mineral tryptone glycerol (MTG) medium, as previously described^53^. We used *nrdA* sequencing to confirm the species assignments of each strain used. For immunostaining analyses, we conjugated pJT04, a vector expressing mCherry with *tetAR* cassette, into AC047Δ*sctV*. We established the parity of cytotoxicity results for tagged and untagged AC047Δ*sctV* (not shown). Additionally, QV306 was conjugated with pJT05, a vector expressing mCherry, for trafficking analysis. All strains used in this work are listed in **Table S2**.

### Mutagenesis

A protocol adapted from *Burkholderia*^42^ was used to generate clean deletions in AC047 and disruption mutants in our *A. xylosoxidans* strains. Briefly, we created suicide vectors containing upstream and downstream regions from our target genes. The vectors had a Sce-I excision site, so that after integration we could introduce an excision vector encoding the restriction enzyme to disrupt the gene. We checked putative mutants by PCR with primers upstream of the US and downstream of the DS, and amplicons of the correct size were sequenced. Suicide and excision vectors were sequentially introduced into *Achromobacter* via conjugation with RHO3, an auxotrophic donor strain requiring diaminopimelic acid^54^. The plasmids used for each mutant are listed below (**Table S2**). We used plasmids derived from *Bordetella* (pDA17 encoding a C-terminal FLAG tag and pDA12 for untagged) for complement expression^55^. Constructs were generated by Gibson assembly^56^, with a few created by restriction/ligation, as designated (**Table S3**).

### In vitro Infection

Bacteria were prepared by 3 washes in RPMI and adjusted by OD to MOIs of 20 and 80, as indicated. Heat-killed bacteria (60°C 20 minutes) served as controls. Cells were infected with bacteria and spun down at 200 xg for 5 minutes to synchronise infection. Cells were incubated at 37°C at various times from 15 minutes up to 5 hours p.i. For transwell experiments, HMDMs were plated in 24-well plates and bacteria were added in the upper chamber in 20% total media volume at the same MOI as during direct infection, following the protocol outlined by Flaherty and Lee^31^.

### LDH assay

Supernatants from infected cells were evaluated using the Roche LDH assay kit. This assay detects lactate dehydrogenase in the medium, which indicates cell membrane rupture. The assay reagents were prepared, and 50 μl were added to 50 μl of each sample media. Data represent four technical replicates per biological replicate, with uninfected and 1% Triton X-100-treated (15 minutes) cells serving as the low (0%) and high (100%) controls respectively. For caspase inhibition, cells were treated with 10 μg/ml Z-VAD-FMK (or equivalent volume DMSO) for an hour pre-infection, and the same concentration during a 3-hour infection.

### Propidium iodide uptake assay

HMDM were plated at a density of 5 x 10^4^ cells per 100ul in transparent-base, black-sided 96 well plates. On the day of infection, 2 ng/ml propidium iodide was added to all test condition wells along with the bacteria. Propidium iodide-free cells were used as a blank, and 1% Triton X-100 cells provided a fluorescence normalisation reading. We took fluorescence readings every 20 minutes using a PolarStar plate reader with 5% CO2 at 37°C. For the internalisation assay, cells were treated with 5 μg/ml cytochalasin D (or equivalent volume DMSO) for an hour pre-infection, and the same concentration during infection.

### Immunoblotting

Lysates were prepared by washing cells with PBS twice, then adding 2% SDS 66mM Tris lysis buffer (10 μl per 100,000 cells) and scraping lysates into Eppendorf tubes on ice. Protein concentration was determined by BCA, and equivalent lysates loaded onto SDS-PAGE gels. We used a Trans-Blot Turbo Transfer then blocked with casein-TBS for 1 hour. Blots were incubated in anti-GSDMD primary antibody in TBS-T, rocking overnight at 4°C, washed thrice with TBS-T, then incubated at room temperature in secondary antibody conjugated to fluorophore for 1 hour. Blots were visualised using a LI-COR system. Blots were stripped and re-probed with anti-GAPDH, a loading control. Antibodies, codes, and concentrations are listed in **Table S1**.

### ELISAs

HMDMs were infected as described above. At five hours post-infection, plates were spun down (5 minutes 200 g), and supernatants were collected and immediately used for analysis. We used the R&D Technologies IL-1β ELISA kit (**Table S1**), following manufacturer instructions and including 4 technical replicates per biological replicate. The mouse cytokine levels were detected in the same way, using kits for IL-6 and KC. Details follow about lung homogenate preparation.

### Immunostaining

UV-sterilised coverslips were placed in 12-well cell culture plates prior to seeding of THP-1 and HMDM. Infections were carried out as described above, and at various time points the coverslips with cells were retrieved, fixed in 4% paraformaldehyde, washed with PBS, and then placed in 14 mM ammonium chloride overnight at 4°C to quench free aldehyde groups. Cells were permeabilised using 0.5% saponin^57^, and then incubated in a humidified chamber at room temperature for an hour with primary antibodies (**Table S1**). Coverslips were washed twice in PBS, then incubated an hour in secondary goat anti-rabbit AlexaFluor 488 antibody, as above. Slides were imaged using SP8 confocal and Stellaris-5 confocal microscopy, images taken at random.

### Live imaging of infected cells

THP-1s were seeded directly into Ibidi 8 well chamber slides. Infections were carried out as described above, at various timepoints the cells were retrieved and incubated with fluid phase markers (**Table S1**). Cells were immediately imaged by Stellaris-5 confocal microscopy.

### Galleria mellonella infections

For each experiment, waxworm larvae were visually inspected and weighed (0.25 to 0.35 g) before injection, to ensure consistency across conditions. We did dose response curves to determine the optimal CFU for injection. Larvae were injected as described ^58^, with 10 μl sterile PBS or 10 μl bacteria of total CFU 10^6^ of wildtype AC047 or AC047Δ*sctV*. Larvae were placed at 37°C and assessed daily for indications of morbidity or mortality.

### Mice infections

C57BL/6 mice, originally purchased from Charles River Laboratories (UK), were bred in house. All animal work was conducted in accordance with the Animals Scientific Procedures Act (1986). The research was ethically reviewed by both the University Animal Welfare and Ethical Review Body (AWERB) and the Northern Ireland Dept of Health. The research was carried out under approved project licenses PPL2807.

Adult female C57BL/6 mice were used in these studies. Mice were given free access to food and water and subjected to 12h light/dark cycle. A log phase culture of *A. insuavis* AC047 and AC047Δ*sctV* was washed and resuspended in sterile endotoxin-free PBS and diluted based on 1 OD6_00nm_ is equal to 1×10^10^ CFU/ml. Prepared inoculums were plated on MTG plates to confirm CFU dose delivered. Groups of n=5 mice were anesthetized and inoculated intranasally with either 10^7^, 10^8^ or 10^9^ CFU of AC047 or 10^8^ AC047Δ*sctV* or PBS (in 20μL). Mice were sacrificed at 24 hours p.i., unless a predefined humane endpoint was reached prior to that. Lungs were collected into 2 ml of PBS and homogenates were serially diluted and plated on MTG agar overnight at 37 °C for quantification of CFUs. Homogenates were also analysed for IL-6 and KC by ELISA (R&D Systems). To directly compare the wild type and mutant, nine-plus-week old female C57BL/6 mice were infected intranasally with 10^7^ of AC047 or AC047Δ*sctV* (n=10 per group). Mice were sacrificed 24h post-infection and n=7 were analysed for bacterial load and IL-6 and KC using ELISA (R&D systems). A further n=3 animals were analysed using histology. The lungs were inflated with paraformaldehyde (PFA), fixed in PFA for 24 hours and dehydrated. The specimens were then embedded in paraffin, cut in 5μm sections, and stained with haematoxylin-eosin. The protocols for histology scoring have been described before, for alveolar wall damage (mean linear intercept)^59^ and septa thickness^60^ and here they are an average of 5 fields of view of 2 sections for each condition at magnification x20.

### Cblaster and statistics

The synteny of *axlG* and *sctA* genes was determined using cblaster v1.3.16^29^. Briefly, we determined the co-occurrence of the genes using a NCBI’s non-redundant protein sequences (nr) database, with the search term “Achromobacter” accessed on 10.12.2022. We used GraphPad Prism v9.4 to generate graphs and perform statistical tests. LDH, CFU and ELISA data were compared by t-tests. Percentage of intracellular bacteria was assessed by Kruskal-Wallis test, and Mantel-Cox for *in vivo* model survival data.

## Acknowledgements

H.P. was supported by a fellowship from the Department of Economy, Northern Ireland. P.Z. was supported by BactiVax, MSCA-ITN-2019, Innovative Training Networks. This research was supported by grant VALVAN19G0 from the Cystic Fibrosis Foundation to M.A.V. R.C.C. was supported by an Academy of Medical Sciences Springboard Award (SBF005\1104), a Royal Society Research Grant (RGS\R1\201127), and a Biotechnology and Biological Sciences Research Council New Investigator Research Grant (BB/V016741/1). C.E.B. received funding from a Wellcome Investigator Award 108045/Z/15/Z and Medical Research Council grant MR/X000826/1. We thank Dr. Janet Torres for the construction of plasmids pJT04 and pJT05, Dr. Annette Vergunst for the gift of pIN62, and Dr. Helina Marshall for technical assistance with the *in vivo* experiments. Strain QV306 was a gift from Prof. M. Tunney, School of Pharmacy, Queen’s University Belfast.

