## Supplemental Figures and Tables for "The *Achromobacter* Type 3 secretion system drives pyroptosis and immunopathology via independent activation of NLRC4 and NLRP3 inflammasomes"

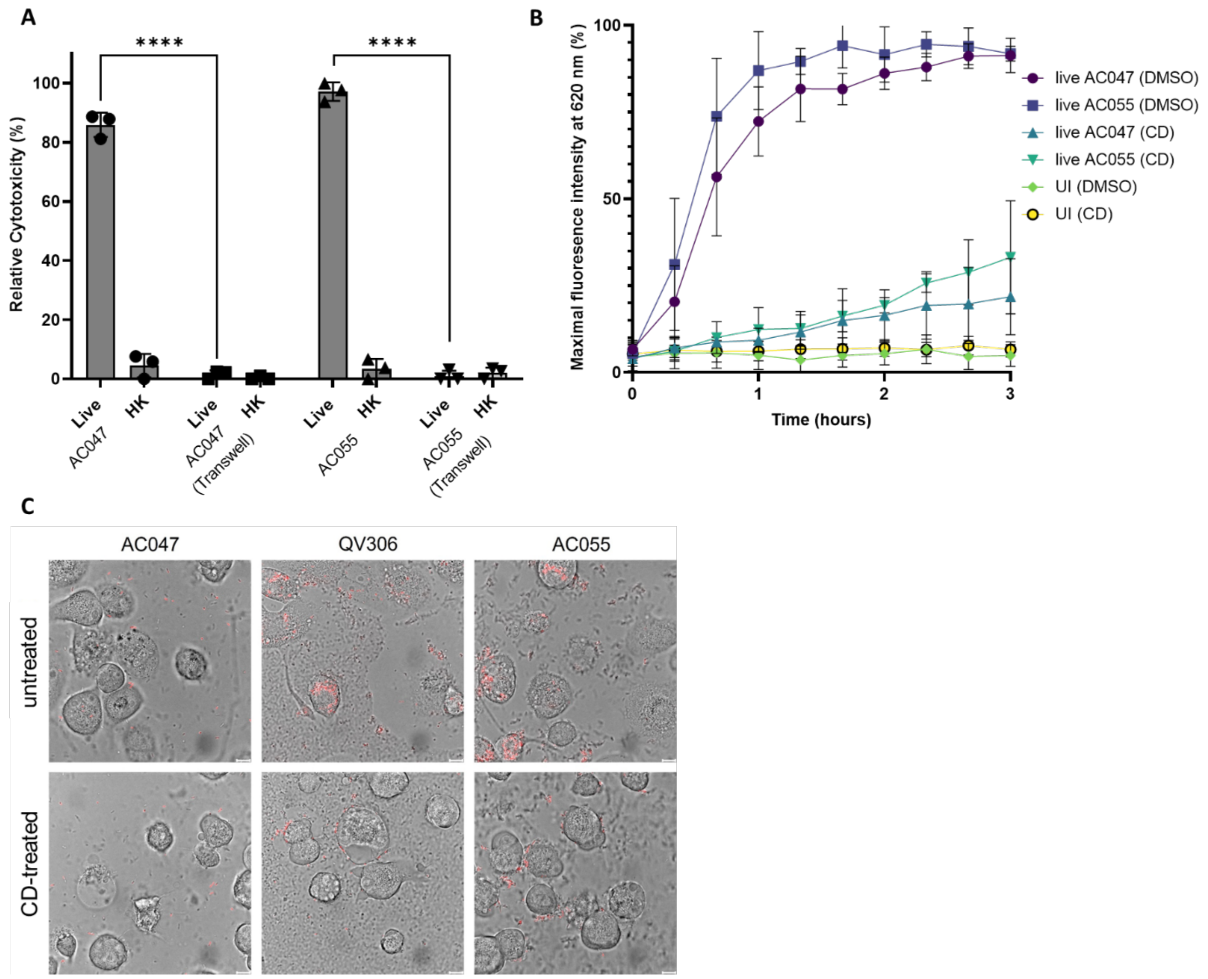

**Figure S1. Cytotoxicity is contact-dependent and enhanced by internalisation**. **A**. LDH release of HMDM infected with live and heat-killed (HK) AC047 and AC055, comparing direct infection and transwell-based at 5 hours p.i., MOI 20 **** signifies p<0.0001 by t-test, error bars SD. **B.** Propidium iodide uptake assay of HMDM, untreated or treated with 5 µg/ml cytochalasin D. Data represent four technical replicates for each of three biological replicates. **C.** THP-1 macrophages 5 hr p.i. with mCherry labelled QV306, AC047 and AC055, untreated or treated with 5 µg/ml cytochalasin D. Images taken with × 100 magnification on Leica Stellaris-5 confocal microscope. MOI = 80 (QV306) and 20 (AC047, AC055). Scale bar = 10 µm. Images taken with × 63 magnification on Leica SP8 confocal microscope. MOI = 20. Scale bar = 10 µm.

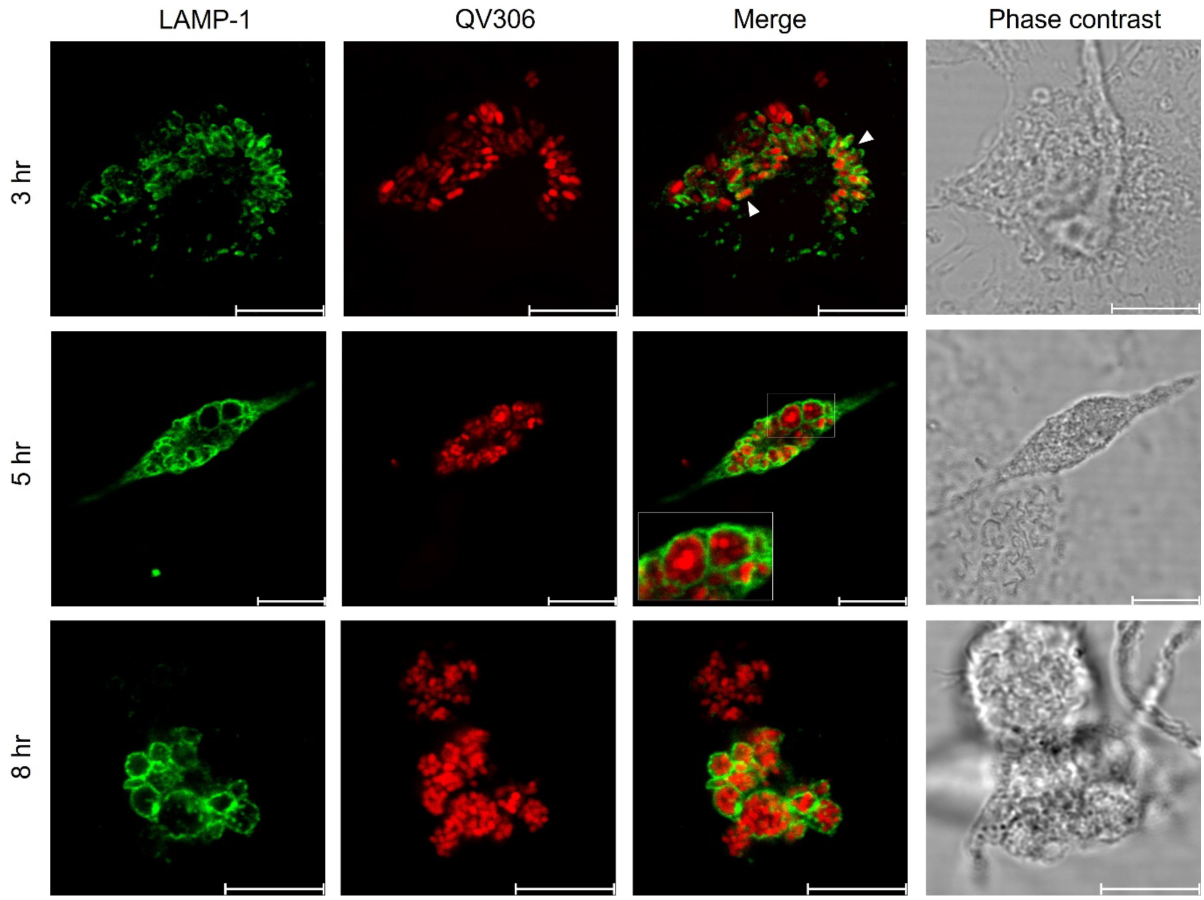

**Figure S2:** AcV containing **QV306 colocalises with LAMP-1 in HMDM.** THP-1 macrophages at 3, 5 and 8 hr P.I with live *A. xylosoxidans* colocalising with LAMP-1. Images taken with × 63 magnification on Leica SP8 confocal microscope. MOI = 80. Scale bar = 10 µm.

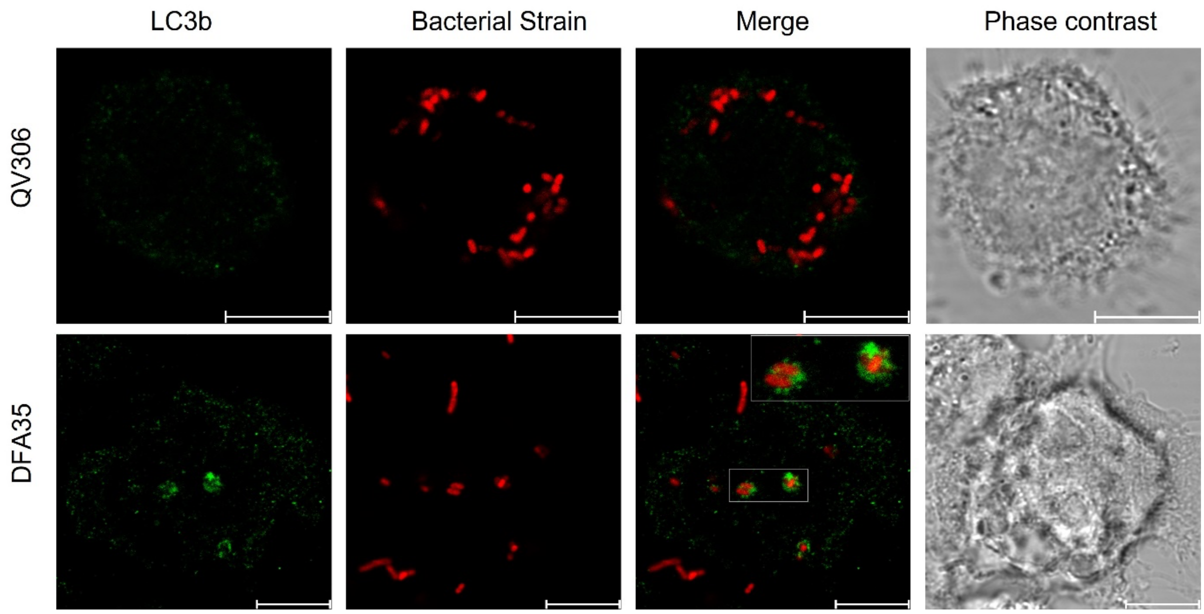

**Figure S3: The QV306 AcV does not have the autophagosome marker LC3b.** THP-1 macrophages 3 hrs P.I immunofluorescently stained with anti-LC3B polyclonal antibody. Positive control is *Burkholderia cenocepacia* isolate DFA35. Images taken with × 63 magnification on Leica SP8 confocal microscope, MOI = 80. Scale bar = 10 µm.

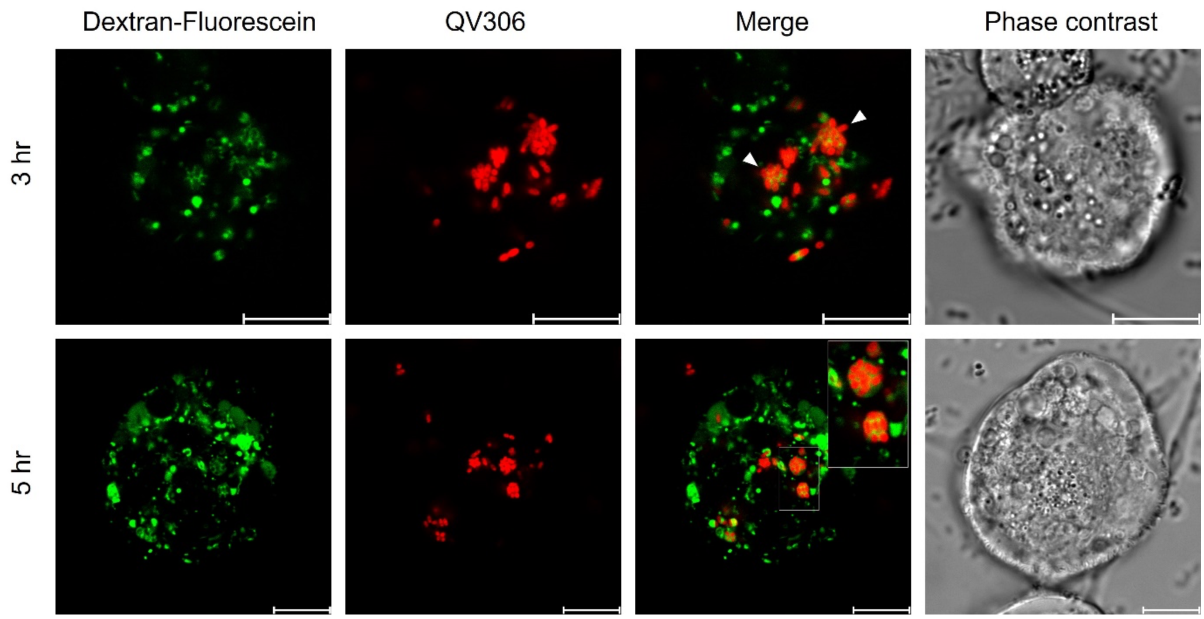

**Figure S4: The QV306 AcV co-localises with Dextran-fluorescein.** THP-1 macrophages 3 and 5 hrs P.I pre-treated with the fluid phase marker dextran fluorescein. Images taken with × 100 magnification on Leica Stellaris-5 confocal microscope. MOI = 80. Scale bar = 10 µm.

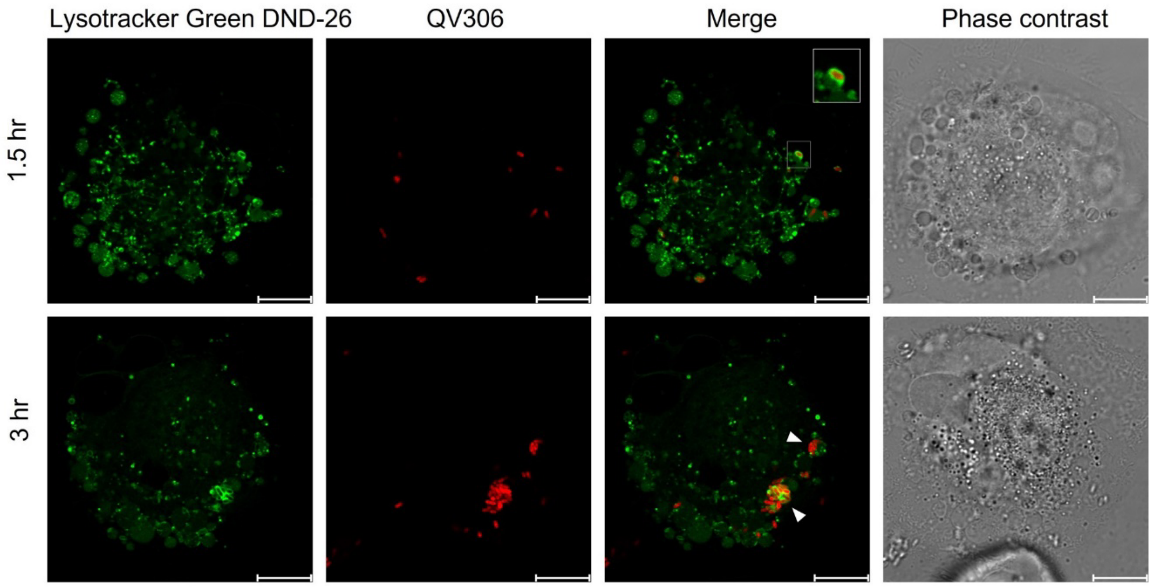

**Figure S5: The QV306 AcV can colocalise in an acidic compartment.** THP-1 macrophages 1.5 and 3 hrs P.I treated with the fluid phase marker Lysotracker green DND and immediately imaged on the Leica Stellaris-5 confocal microscope, × 100 magnification. MOI = 80. Scale bar = 10 µm.

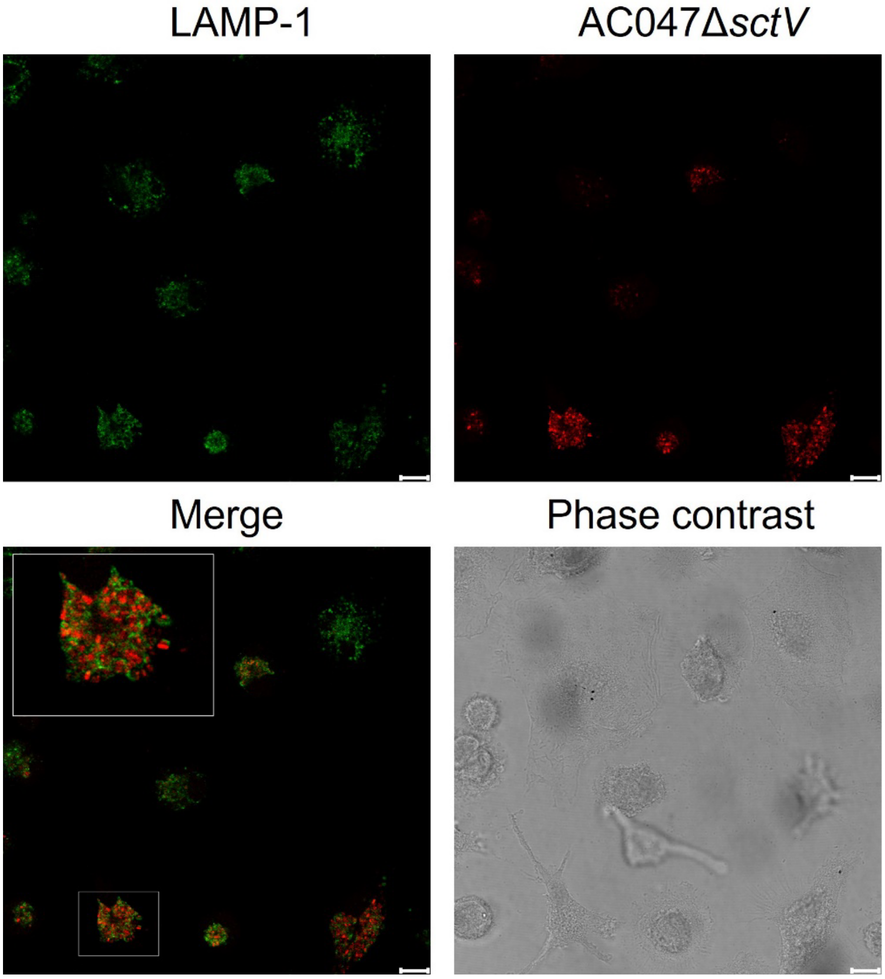

**Figure S6. mCherry-labelled AC047ΔSctV 24 hours post-infection of HM**

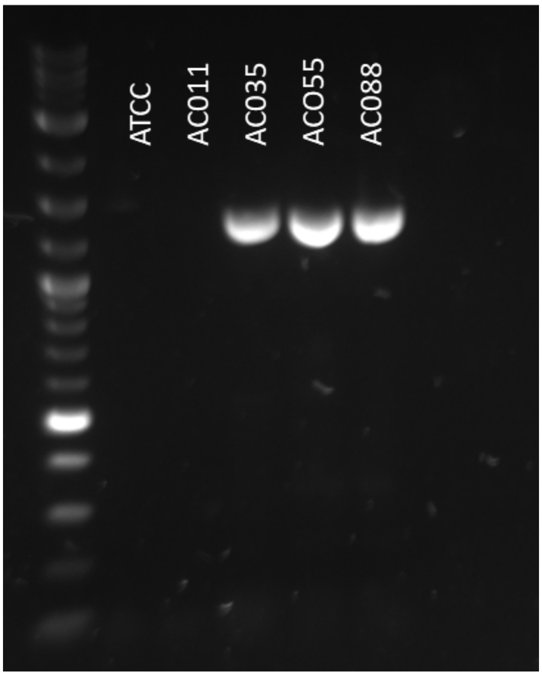

**Figure S7:** PCR of *sctN* (T3SS component) in various *Achromobacter* strains.

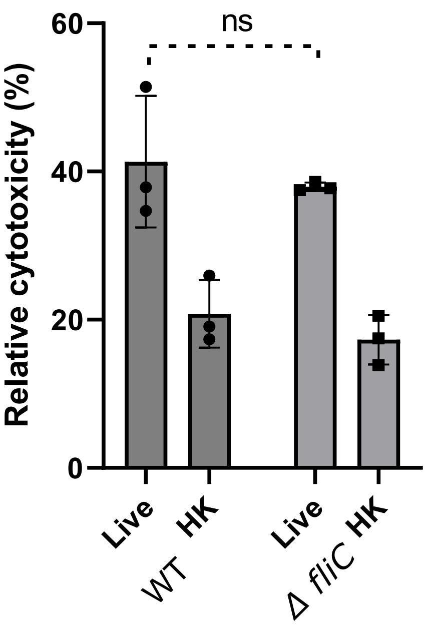

**Figure S8.** LDH release of NLRC4 KO THP-1 infected with WT or Δ*fliC* AC047. Data represents three biological replicates.

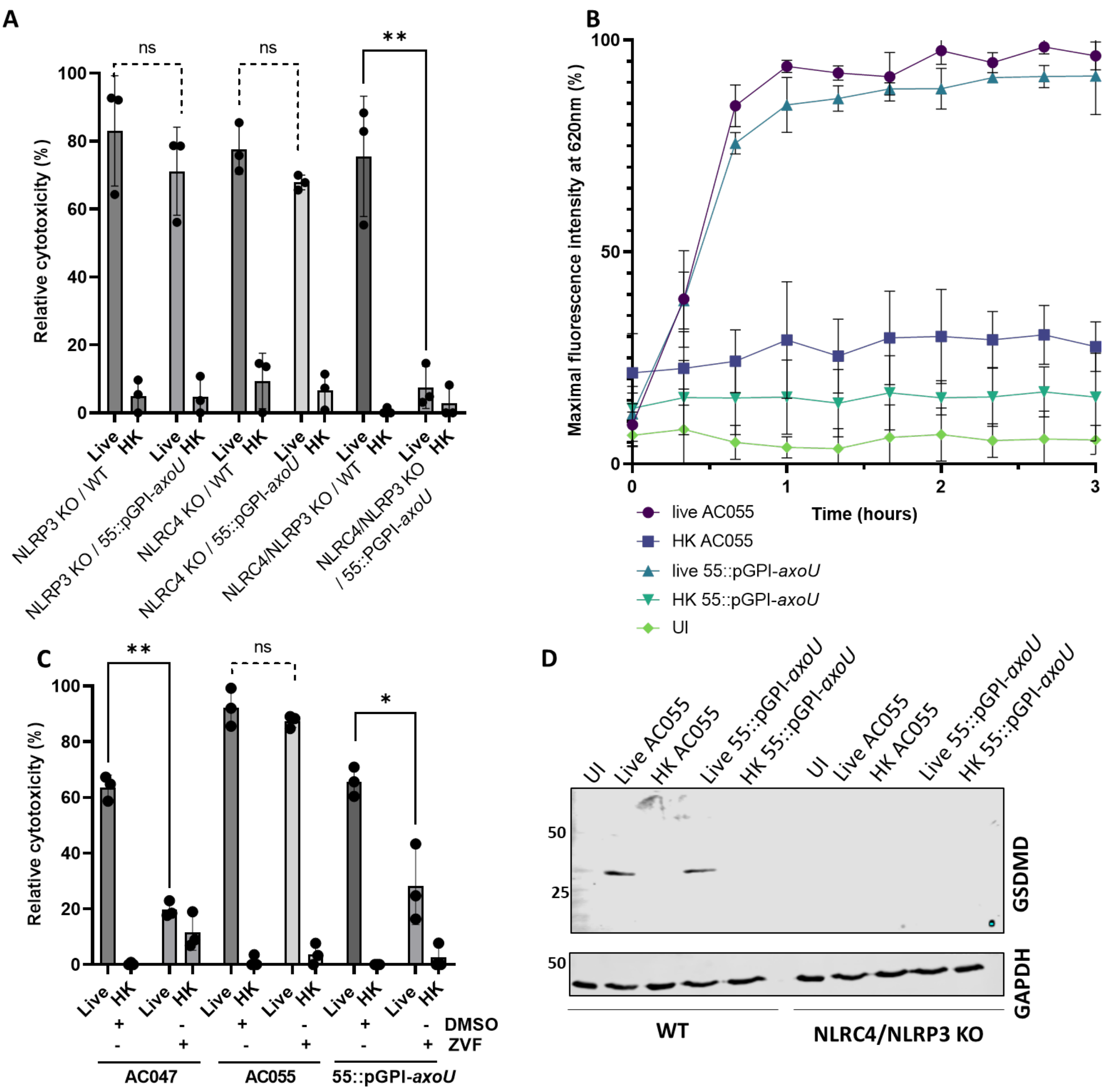

**Figure S9. AxoU is not necessary or sufficient for pyroptosis. A.** LDH assays of AC055 and disrupted AxoU mutant infecting THP-1 KO cells. **B.** PI assay cytotoxicity time-course of HMDM infected with AC055 or with AC055::pGPI-*axoU*. **C.** LDH assay of HMDM cells infected with live/HK bacteria and treated with 10 µg/ml Z-VAD-FMK or equivalent volume DMSO, 3-hour infection. **D.** AC055::pGPI-*axoU* induces pyroptosis in WT THP-1, but neither wildtype AC055 nor AC055::pGPI-*axoU* can induce Gasdermin-D cleavage in dKO THP-1. ** signifies $p<0.005$, * $p<0.05$ by t-test, MOI 20, error bars SD. Data represents three biological replicates.

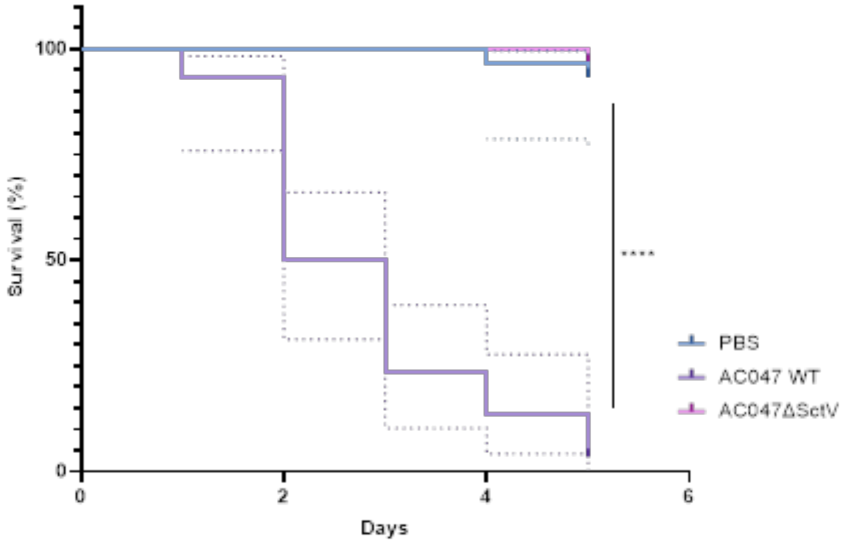

**Figure S10. AC047 requires a functional T3SS to efficiently kill *Galleria mellonella.*** Larvae were injected with $10^6$ CFU (10 µl), PBS was used as a control. Data shown are representative of 3 independent replicates, each with 10 waxworms per test condition. **** signifies p<0.0001 by Mantel-Cox test.

**Table S1. Reagents/Resources**

| Reagent or Resource | Source | Identifier | Notes/reference |
|---|---|---|---|
| ***Antibodies*** | | | |
| Gasdermin-D | Cell Signalling Technology | #93709 | 1:1000 |
| Cleaved Gasdermin-D (N-terminus) | Cell Signalling Technology | #36425 | 1:1000 |
| GAPDH | Abcam | ab8245 | 1:5000 |
| EEA1 | Invitrogen | PA1-063A | 1:1142 |
| LAMP-1 | Abcam | ab24170 | 1:800 |
| LCB3 | Invitrogen | L10382 | 0.5 µg/ml |
| (IgG control) | Abcam | ab171870 | (Equivalent µg/ml to LAMP-1 ab) |
| Rabbit IgG | LI-COR | 925-32211 | 1:10,000 |
| Mouse IgG | LI-COR | 926-32210 | 1:10,000 |
| Rabbit IgG (Alexa488) | Abcam | ab150077 | 1:2000 |
| ***Cells and culture reagents*** | | | |
| THP-1 | American Type Culture Collection | TIB-202™ | |
| NLRP3 KO THP-1 | | | [52] |
| NLRC4 KO THP-1 | | | [37] |
| NLRC4/NLRP3 KO THP-1 | | | [37] |
| Peripheral blood buffy coats | Northern Ireland Blood Transfusion Service | | |
| RPMI 1640 Medium | ThermoFisher Scientific | 11875093 | |
| Fetal Bovine Serum, qualified, heat inactivated, Brazil | ThermoFisher Scientific | 10500064 | |
| Penicillin-Streptomycin (10,000 U/mL) | ThermoFisher Scientific | 15140122 | |
| IMDM | ThermoFisher Scientific | 12440053 | |
| Phorbol 12-myristate 13-acetate | Sigma-Aldrich | P8139-1MG | |
| Recombinant Human GM-CSF | Peprotech | 300-03 | |
| ***Chemicals*** | | | |
| Agar | Melford | A20250 | |
| Ammonium chloride | Biosciences | #RC-015 | |
| Ammonium sulfate | Honeywell | A5132-1KG | |
| CD14 Microbeads 2ML | Miltenyi Biotec | 130-050-201 | |
| Chloramphenicol | Sigma-Aldrich | C0378-25G | |
| Cytochalasin D | Sigma-Aldrich | C2618-200UL | 5 µg/ml |
| Dextran | ThermoFisher Scientific | D1821 | 50 µg/ml |
| DMSO | Sigma-Aldrich | D8418 | |

| | | | |
|---|---|---|---|
| Fluoroshield with 1,4-Diazabicyclo[2.2.2]octane | Sigma-Aldrich | F6937-20ML | |
| Glycerol | Sigma-Aldrich | G5516 | |
| High salt LB | Melford | L24040 | |
| Lysotracker | Invitrogen | L7526 | 250 nM |
| Paraformaldehyde Solution, 4% in PBS | ThermoFisher Scientific | 15670799 | |
| Phosphate buffered saline | Sigma-Aldrich | P4417-100TAB | |
| Potassium phosphate monobasic | Sigma-Aldrich | P5379-500G | |
| Potassium nitrate | Sigma-Aldrich | P8291-KG | |
| Propidium iodide | Sigma-Aldrich | P4864-10ML | |
| Sodium dihydrogen orthophosphate dihydrate | GPR | 301324Q | |
| Tetracycline hydrochloride | Sigma-Aldrich | T3383-100G | |
| Triton X-100 | Bio-Rad | 1610407 | |
| Tryptone | Sigma-Aldrich | T9410-1KG | |
| Tween-20 | Sigma-Aldrich | P1379-500ml | |
| Z-VAD-FMK | InvivoGen | tlrl-vad | 10 µg/ml |
| ***Critical commercial assays, enzymes, and kits*** | | | |
| ROCHE Cytotoxicity Detection Kit (LDH) | Scientific Laboratory Supplies | 11644793001D2 | |
| Human IL-1 beta/IL-1F2 DuoSet ELISA | R&D Systems | DY201-05 | |
| Mouse CXCL1/KC DuoSet ELISA | R&D Systems | DY453-05 | |
| Mouse IL-6 DuoSet ELISA | R&D Systems | DY406-05 | |
| DuoSet ELISA Ancillary Reagent Kit 2 | R&D Systems | DY008B | |
| Q5® Hot Start High-Fidelity DNA Polymerase | New England BioLabs | M0493S | |
| Trans-Blot Turbo RTA Mini 0.2 µm Nitrocellulose Transfer Kit, for 40 blots | Bio-Rad | 1704270 | |
| ***Experimental models: Organisms/strains*** | | | |
| C57BL/6 (mouse) | | | |
| ***Software*** | | | |
| GraphPad Prism v. 9.4 | GraphPad Software | | |
| Benchling | Benchling [Biology Software]. (2022) | | |
| ***Other*** | | | |
| Transwells | Sarstedt | 83.3932.040 | |
| LS Columns 25/PK | Miltenyi Biotec | 130-042-401 | |
| Ficoll® Paque Plus | Sigma-Aldrich | 17-1440-03 | |
| PolarStar plate reader | BMG Labtech | 415-201 | |

| | | |
|---|---|---|
| Trans-Blot Turbo Transfer Starter System | Bio-Rad | 17001918 |
| SP8 confocal microscope | Leica | |

**Table S2. Strains and Plasmids**

| ***Strain*** | ***Relevant characteristics*** | ***Source and/or reference*** |
|---|---|---|
| *Achromobacter* | | |
| AC011 | *A. xylosoxidans* | |
| AC035 | *Achromobacter* sp. | |
| AC047 | *A. insuavis* | |
| AC055 | *A. xylosoxidans* | |
| AC088 | *A. xylosoxidans* | [53] |
| ATCC 27061 | *A. xylosoxidans* | American Type Culture Collection |
| QV306 | *A. xylosoxidans* | Michael Tunney |
| *Burkholderia* | | |
| DFA35 | *B. cenocepacia* Δ*atsR* | [61] |
| *Escherichia coli* | | |
| DH5α | F - ϕ80*lacZ* M15 *endA1 recA1 supE44 hsdR17*($r_K^-$ $m_K^+$) *deoR thi-1 nupG supE44 gyrA96 relA1* Δ(*lacZYA-argF*)U169, λ– | Laboratory stock |
| RHO3 | *Δasd thi-1 thr-1 leuB26 tonA21 lacY1 supE44 recA; integrated RP4-2 Tcr::Mu ΔaphA* (*λ*$pir^+$) | [54] |
| SY327λ*pir* | *F-, araD, Δ(lac pro), argE(Am), recA56,* $Rif^R$, *gyrA λpir* | [62] |
| ***Plasmid*** | | |
| pGPI-SceI-XCm | Suicide plasmid vector, R6Kɣ origin of replication, $Mob^+$, carries a I-SceI endonuclease site; $Cm^R$ | [42] |
| pGPI-SceI-XCm-*sctV*(Ax) | *sctV*(Ax) mutagenic plasmid | This study |
| pGPI-SceI-XCm-*sctV*(Ai) | *sctV*(Ai) mutagenic plasmid | This study |
| pGPI-SceI-XCm-*sctX* | *sctX* mutagenic plasmid | This study |
| pGPI-SceI-XCm-*axlG* | *axlG* mutagenic plasmid | This study |
| pGPI-SceI-XCm-*hcp* | *hcp* mutagenic plasmid | This study |
| pGPI-SceI-XCm-*axoU* | *axoU* mutagenic plasmid | This study |
| pDAI-SceI-sacB | Expresses the I-SceI endonuclease, *sacB* gene; $Tet^R$ | [42] |
| pDA12 | Expression vector (untagged); $Tet^R$ | [63] |
| pDA12-*sctX* | pDA12 carrying *sctX* | This study |
| pDA17 | Expression vector to construct C-terminal FLAG-tagged protein fusions; $Tet^R$ | [63] |
| pDA17-*axlG* | pDA17 carrying $axlG_{FLAG}$ | This study |
| pJT04 | pDA12 encoding the mCherry red fluorescent protein; $Tet^R$ | J. Torres Bustos |
| pIN62 | $ori_{pBBR}$ $mob^+$, $Cm^R$, DSRed gene | [64] |
| pJT05 | pIN62, DSred gene replaced by the mCherry red fluorescent protein gene; $Cm^R$ | J. Torres Bustos |

### Table S3. Primers

| Gene | Use[a] | Forward primer[b] | Reverse primer[b] |
|---|---|---|---|
| *nrdA* | **SS** | GAA CTG GAT TCC CGA CCT GTT | TTC GAT TTG ACG TAC AAG TTC TGG |
| *sctN* | **SS** | ATGCGCCAGTTCGACTACATCTCCGAAATGATG | GCCGCCCCCACGATGGCG |
| ***A. insuavis*** | | | |
| *sctV* | **US** | cgtgctgacctgacctgagcGCCACATCATCGAGACCTTC | gatgttgtagCCGATGGTCAGGATGGAG |
| | **DS** | tgaccatcggCTACAACATCCTGCTGTCCGAAG | cgacggatcccaagcttcttAGCAGGTAAGCGGGCAGG |
| | **SS** | CACCATCGACATGGACGAGGCG | TGCAGGTTCTGTTCCGTGTCGC |
| *axlG* | **US** | cgtgctgacctgacctgagcCATGCAGGCCAAGCTCAAC | gtgcgcggacCCATTGCCGGTCTCCGGA |
| | **DS** | ccggcaatggGTCCGCGCACGATGGTCTGAC | cttctagacggtaccgcatgAGCGGGTCGAGCAGCGCG |
| | **SS** | ATCGAGGAAGTGCAGAAGC | TTCCAGCAGGCATTGGG |
| | **Comp** | gaaggattcggaattctacaATGGCCGACACCCAAGACC | cgtcgtcgtccttgtagtctACCATCGTGCGCGGACG |
| *sctX* | **US** | cgtgctgacctgacctgagcACCATTCGCATCTGGCGG | gttcatccctGCGCCAGTTCCTGGCGTT |
| | **DS** | gaactggcgcAGGGATGAACGCCTCCATGCCC | cttctagacggtaccgcatgCGGGGCGCAGGGCAACGT |
| | **SS** | ACGAAGTGCTGGTCTACG | CATCATGAAGATGATCGAGACG |
| | **Comp** | gaaggattcggaattctacaATGTCCGATATGCGCATCAG | tgcatgcctgcaggtcgactTCATCCCTGGTACAGCGC |
| *hcp* | **US** | AGACGCGCATGCGCGCAGTCGGTGCAGAAGG | TTGGTCGGTACCGTCCGGGTGATCCTTGTCCTGG |
| | **DS** | AACCAGGGTACCTGGAGCCTGACCAAGAACG | AGACGCTCTAGAGGTCGCGTCGAACGTCACC |
| | **SS** | TGGTCGAGAACCTGCCGA | TGACCGAATATTGCAGCG |
| ***A. xylosoxidans*** | | | |
| *axoU* | **MS** | cgtgctgacctgacctgagcACGCGGCGACAAGTTCGTG | cttctagacggtaccgcatgGGGCGAGTTGCCGGTCGT |
| | **SS** | ccagttcgatctgctgttcgacgacc | gcggaagacctgctgcgttcg |
| *sctV* | **US** | TGCTGGCGCGGCCGCGGCCACATCATCGAGACCTTCGG | CGTCGCAGCATGCGCCGTCGCCGATGGTCAGGAT |
| | **DS** | GCTGGAGGCATGCCTACAACATCCTGCTGTCCGA | GCCACCATCTAGACGCGCTTGATGTTGTCCACCAGCTT |
| | **SS** | caccatcgacatggacgaggcg | tgcaggttctgttccgtgtcgc |

[a] US = upstream, DS = downstream, SS = Sanger sequencing, MS = mid-sequence, Comp = complement.

[b] Lower case denotes vector overlap sequence for Gibson assembly, restriction sites are underlined. *A. xylosoxidans* mutants were gene disruptions, so we used the primers that recognised our suicide vector in combination with listed SS primers. All sequences given in 5' to 3'.
